# ContrasTED: contrastive domain embeddings for scalable remote homology classification

**DOI:** 10.64898/2026.08.28.747864

**Authors:** David Miller, Nicola Bordin, Janusan Jeyananthan, Vaishali Waman, Michael Heinzinger, Christine Orengo

**Affiliations:** Institute of Structural and Molecular Biology, University College London, London, UK; Centre for Artificial Intelligence, University College London, London, UK; Department of Biomedical Sciences, University of Padova, Italy; Institute of Computational Biology, Computational Health Center, Helmholtz Munich, Germany; School of Computation, Information and Technology, Technical University of Munich, Germany

**Keywords:** protein domains, homology, contrastive learning, protein language models

## Abstract

Protein structure prediction has expanded structural databases to hundreds of millions of domains. Classifying these domains into homologous superfamilies reveals evolutionary and functional relationships that can persist despite low sequence similarity. As the size of structural databases continues to grow, homology classification requires methods that combine scalability with accuracy. Here we present ContrasTED, which uses CATH-supervised center-contrastive learning to project structure-aware embeddings into a domain-level metric space for nearest-centroid superfamily assignment. On a sequence-filtered S20 benchmark (*n* = 1,028), superfamily assignment accuracy reached 92.9% (1-NN) and 91.4% (nearest centroid), exceeding sequence search, profile HMMs, Foldseek, and a classifier trained on embeddings. The learned latent space separates superfamilies while retaining structural information below 20% sequence identity, with the largest gains among sparsely represented superfamilies. ContrasTED produces 4.67 million new candidate assignments across 3,796 superfamilies in The Encyclopedia of Domains (TED), extending annotation coverage beyond previous structure-based methods.

## 1 Introduction

The AlphaFold Database contains more than 214 million predicted structures, which have been divided into approximately 365 million domains [14, 16, 17, 30, 32]. Resources such as the Encyclopedia of Domains (TED) organize this expansion by segmenting predicted structures into domains and placing them within structural classifications [16]. Their scale creates a general need for methods that can classify millions of domains while retaining sensitivity to remote evolutionary relationships.

Hierarchical classifications such as CATH organize domains by Class (C), Architecture (A), Topology (T), and Homologous Superfamily (H) [22, 31]. Assignment to the H level indicates shared ancestry, providing a reliable foundation for transferring evolutionary and functional hypotheses. Detecting homology from sequence alone breaks down in the sequence twilight zone below 20–30% pairwise identity, where alignment scores become indistinguishable from background noise [1, 25]. Profile HMMs expand this boundary by capturing position-specific conservation and underpin large-scale resources such as Gene3D, Pfam, and TEDLH [2, 5, 18, 19], yet searching massive profile libraries remains computationally intensive. Because 3D structure is more conserved during evolution than sequence [13], structural comparison provides a powerful route to identify remote homologues. However, classical structural alignment algorithms such as TM-align, SSAP and DALI are far too computationally expensive to run all-against-all across millions of structures [12, 23, 34]. Fast structural search tools, including Foldseek and Foldclass, address this computational bottleneck by converting 3D backbones into discrete structural alphabets or geometric embeddings, making large-scale structural comparison practical [15, 29].

Alongside sequence and structural alignment, pretrained protein language models (pLMs) provide rich representations for remote-homology detection. ProtT5 learns sequence representations through self-supervised pretraining on millions of protein sequences, while ProstT5 extends this framework by jointly encoding amino-acid sequences and Foldseek 3Di structural tokens [6, 10]. These representations can be transferred directly via nearest-neighbour lookup [26], used to predict structural similarity for remote-homology search (TM-Vec) [8], aligned dynamically at the per-residue level (EBA) [24], or adapted using supervised classifiers such as CATHe and CATHe2 [20, 21]. To optimize embeddings specifically for structural retrieval, ProtTucker applied contrastive learning to CATH using a hierarchy-aware triplet loss [9]. However, pair- and triplet-based contrastive learning requires at least two members of a superfamily to co-occur within the same mini-batch to form a valid positive pair. This constraint is restrictive in CATH, where superfamily sizes are heavily skewed: a small number of superfamilies contain many structures, but most are represented by only one or two domains, making it difficult for rare superfamilies to participate in batch updates.

Here, we present ContrasTED, a structure-aware metric-learning method for remote-homology classification across CATH. ContrasTED concatenates amino-acid and 3Di embeddings from ProstT5 and projects them into a 128-dimensional embedding space using a center-contrastive loss [3]. This proxy-based objective represents each homologous super-family by a learnable center and compares every sampled domain against the entire bank of superfamily centers. As a result, domains from rare or singleton superfamilies contribute direct gradient signals without requiring co-occurring positive pairs in the batch. The model is trained on curated CATH superfamilies augmented with 11.1 million high-confidence TED domains. ContrasTED substantially improves remote-homology classification accuracy over fast structure search tools such as Foldseek and Foldclass on sequence-filtered benchmarks. At inference, query embeddings can be classified either by nearest-neighbour retrieval against individual reference domains or by nearest-centroid retrieval against precalculated super-family means (Fig. 1). While nearest-neighbour lookup evaluates every reference domain, nearest-centroid classification collapses each superfamily into a single normalized prototype, substantially reducing memory and search costs while achieving comparable accuracy. A calibrated distance threshold ensures that accepted queries receive high-confidence superfamily assignments while rejecting ambiguous or out-of-distribution structures. ContrasTED complements residue-level tools such as profile HMMs and EBA by providing rapid, accurate classification across large structural databases.

**Figure 1:**
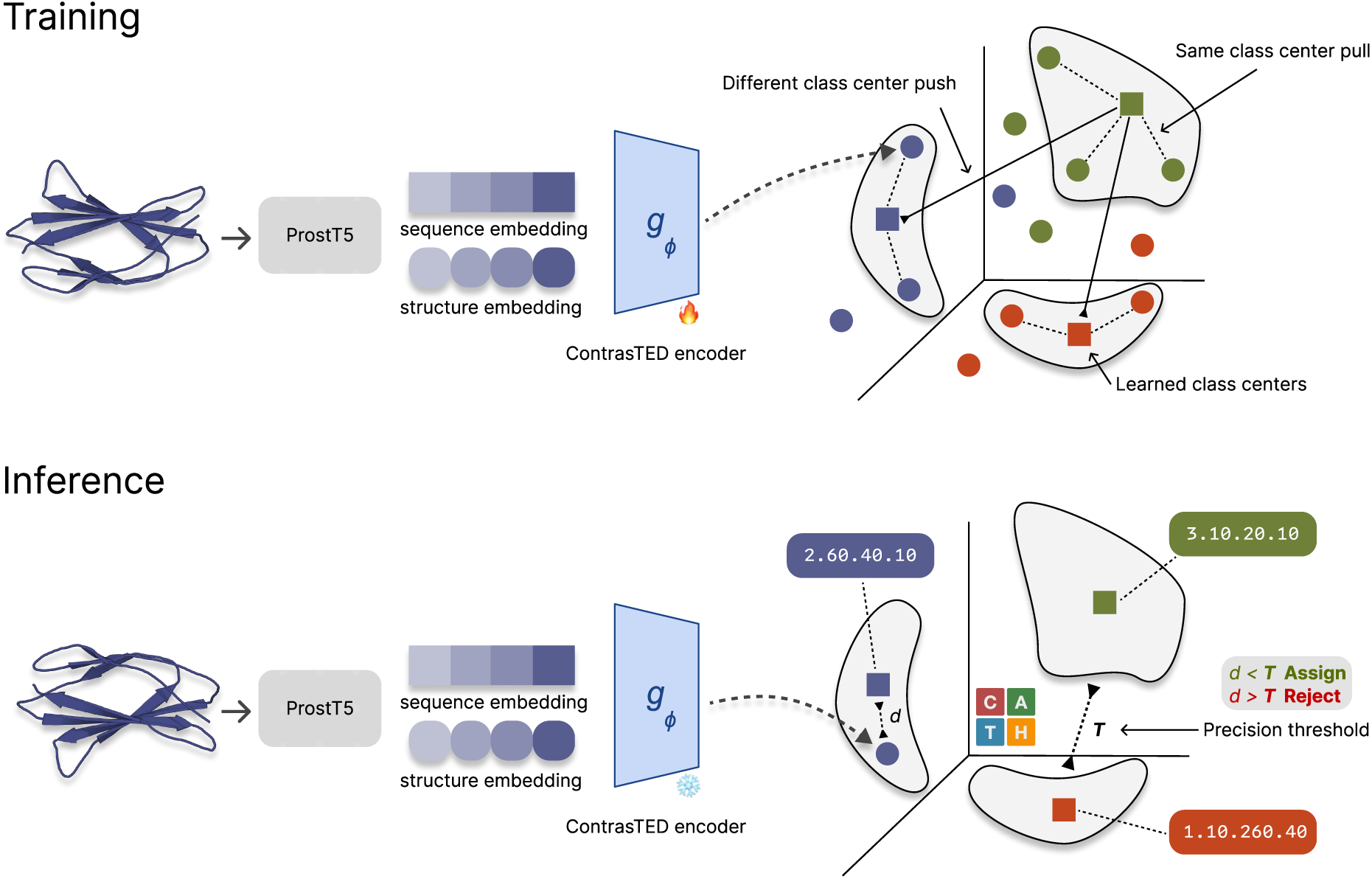
ContrasTED training and inference for remote-homology detection and classification. **Training:** ProstT5 separately encodes the amino-acid sequence and its structure-derived Foldseek 3Di sequence; the mean-pooled embeddings are concatenated and projected by the trainable ContrasTED encoder *g_ϕ_*. Center-contrastive learning pulls each domain towards the learnable proxy center for its CATH H-level superfamily and pushes it away from competing centers, using curated CATH labels together with automated TED labels. **Inference:** after training, *g_ϕ_* is frozen and centroids are calculated from projected CATH reference domains; the learned training centers are not used for retrieval. A query receives the label of its nearest H-level centroid when the cosine distance *d* is below the validation-selected precision threshold *T* and is otherwise rejected.

## 2 Results

### 2.1 ContrasTED improves remote-homology classification

We evaluated ContrasTED on the sequence-filtered S20 benchmark (*n* = 1,028) against sequence search (MMseqs2) [28], profile HMM search (HH-suite) [27], frozen protein language model embeddings (ProtT5, ProstT5), a supervised neural network classifier (CATHe2), contrastive baselines (ProtTucker), and structure-based methods (Foldseek, Foldclass, Progres) [7, 15, 29] (Table 1). Each test set contains exactly one representative domain per superfamily. Accuracy was calculated across all test queries; queries for which a method returned no label were treated as incorrect, making top-1 accuracy equivalent to macro recall across superfamilies.

**Table 1:** Remote-homology CATH assignment accuracy (%). Accuracy on the S20 one-per-superfamily remote-homology test set (*n* = 1,028). Bold marks ContrasTED-AA ∥ 3Di (1-NN and nearest centroid). *^†^*HH-suite3 searches a TEDLH subset restricted to profiles whose CATH seed domain belongs to the training split. This prevents a query from matching a profile seeded on itself and removes profiles seeded on domains withheld from training by the S20 identity filter, matching the CATH-train lookup used by the other methods (profile coverage 93.3%, *n* = 959 of 1,028 queries reachable; on this reachable set, HH-suite3 achieves 89.1% C, 82.8% A, 79.6% T, and 77.5% [74.9–80.1] H accuracy; Supplementary Tables S8–S9). *^‡^*Foldseek 1-NN is by bitscore against the CATH training set. *^§^*CATHe2 is retrained on the matched split and modalities. *^∗^*ContrasTED-AA ∥ 3Di is the mean of seeds 40–42. Confidence intervals and operating details are in Supplementary Tables S1, S6, and S8–S9 and Methods.

| | | Accuracy (%; S20 $n = 1028$) | | | |
| --- | --- | --- | --- | --- | --- |
| Method | Mode | C | A | T | H |
| *Sequence* |  |  |  |  |  |
| MMseqs2 | 1-NN | 39.5 | 37.9 | 37.3 | 36.3 [33.3–39.2] |
| HH-suite3<sup>†</sup> | profile | 85.7 | 78.6 | 74.7 | 72.3 [69.7–74.9] |
| *Structure* |  |  |  |  |  |
| Foldseek (default)<sup>‡</sup> | 1-NN | 94.5 | 92.0 | 89.3 | 85.3 [83.2–87.5] |
| Foldseek (sensitive) | 1-NN | 96.5 | 94.5 | 92.5 | 88.2 [86.2–90.3] |
| Progres | 1-NN | 95.7 | 81.2 | 69.5 | 54.4 [51.5–57.4] |
| Foldclass | 1-NN | 96.5 | 85.1 | 72.8 | 56.8 [53.8–59.9] |
| *Frozen embeddings* |  |  |  |  |  |
| ProtT5 | 1-NN | 93.0 | 83.9 | 75.5 | 69.6 [66.8–72.3] |
| ProstT5-AA | 1-NN | 96.3 | 90.3 | 85.3 | 78.5 [76.1–81.0] |
|  | centroid | 94.8 | 85.2 | 77.3 | 66.8 [63.9–69.6] |
| ProstT5-AA\|\|3Di | 1-NN | 97.6 | 92.1 | 88.4 | 81.9 [79.6–84.3] |
|  | centroid | 96.0 | 89.7 | 83.9 | 73.2 [70.6–76.0] |
| *Supervised ANN* |  |  |  |  |  |
| CATHe2-AA<sup>§</sup> | softmax | 96.5 | 93.0 | 87.6 | 82.1 [80.0–84.5] |
| CATHe2-AA\|\|3Di<sup>§</sup> | softmax | 98.2 | 95.3 | 91.7 | 84.2 [82.1–86.5] |
| *Contrastively projected* |  |  |  |  |  |
| ProtTucker | 1-NN | 95.1 | 88.2 | 82.0 | 75.0 [72.4–77.6] |
| ContrasTED-AA | 1-NN | 97.2 | 93.5 | 91.1 | 88.7 [86.9–90.7] |
|  | centroid | 97.0 | 92.8 | 90.2 | 87.5 [85.4–89.5] |
| ContrasTED-AA\|\|3Di* | 1-NN | **98.9** | **97.3** | **95.9** | **92.9** [91.4–94.3] |
|  | centroid | **98.3** | **96.5** | **95.1** | **91.4** [89.9–93.0] |

Before contrastive projection, incorporating local structural information into the embedding representation provided immediate gains. Concatenating Foldseek 3Di structural tokens with ProstT5 amino-acid embeddings (AA ∥ 3Di) increased frozen 1-NN H-level accuracy from 78.5% for amino acids alone to 81.9%. Adding structural context to the language model embeddings thus brought frozen retrieval close to default Foldseek, which achieved 85.3% across all test queries (88.2% in sensitive mode). HH-suite search of a train-only TEDLH profile subset improved on the sequence-only MMseqs2 baseline of 36.3%, reaching 72.3% accuracy across the full test set and 77.5% on the 93.3% of queries whose superfamily is represented in that subset (Table 1).

Training the projection head with center-contrastive loss and expanding the training data with TED structural representatives yielded substantial additional improvements. Augmenting CATH supervision with 11.1 million TED domains increased 1-NN accuracy to 92.9% (an 11.0 percentage point gain over frozen 1-NN retrieval). Nearest-centroid classification with the complete ContrasTED-AA ∥ 3Di model reached 91.4%, an 18.2 percentage point improvement over frozen nearest-centroid retrieval (73.2%) and 9.5 percentage points above frozen 1-NN retrieval. Nearest-neighbour and nearest-centroid retrieval with ContrasTED yielded closely matched performance across all CATH hierarchy levels (92.9% vs. 91.4% at H-level), demonstrating that each superfamily can be effectively represented by a single normalized centroid vector without substantial loss of classification accuracy.

Sequence-only ContrasTED-AA reached 88.7% 1-NN accuracy, matching Foldseek (88.2% in sensitive mode) despite using no structural input. Among supervised neural networks, CATHe2 retrained on the same CATH split and multimodal inputs achieved 84.2% H-level accuracy. ProtTucker also improved over its frozen ProtT5 baseline, reaching 75.0% accuracy. Structure retrieval methods such as Foldclass and Progres were designed primarily for fold-and topology-level (T-level) retrieval; while they demonstrate high Class and Architecture accuracy, they show lower discrimination at the homologous superfamily (H) level (Table 1 and Supplementary Table S6).

### 2.2 The learned metric increases held-out separation while preserving structural ordering

We first examined how metric learning alters the geometry of held-out superfamily separation. On the S20 test set, we evaluated the centroid decision margin for each query—defined as the distance to the nearest competing centroid minus the distance to the true centroid (*d*_false_*−d*_true_), where positive values indicate a correct top-1 assignment. Following contrastive learning, the number of queries with a positive margin increased from 753/1,028 to 932/1,028, and the median paired margin increased by 0.289 (95% bootstrap CI [0.280, 0.297]; Fig. 2c). Analysis of individual query transitions showed that contrastive learning rescued 190 previously misclassified domains while causing only 11 regressions from correct to incorrect labels (Fig. 2f), demonstrating that the accuracy gains reflect genuine expansion of held-out decision margins.

**Figure 2:**
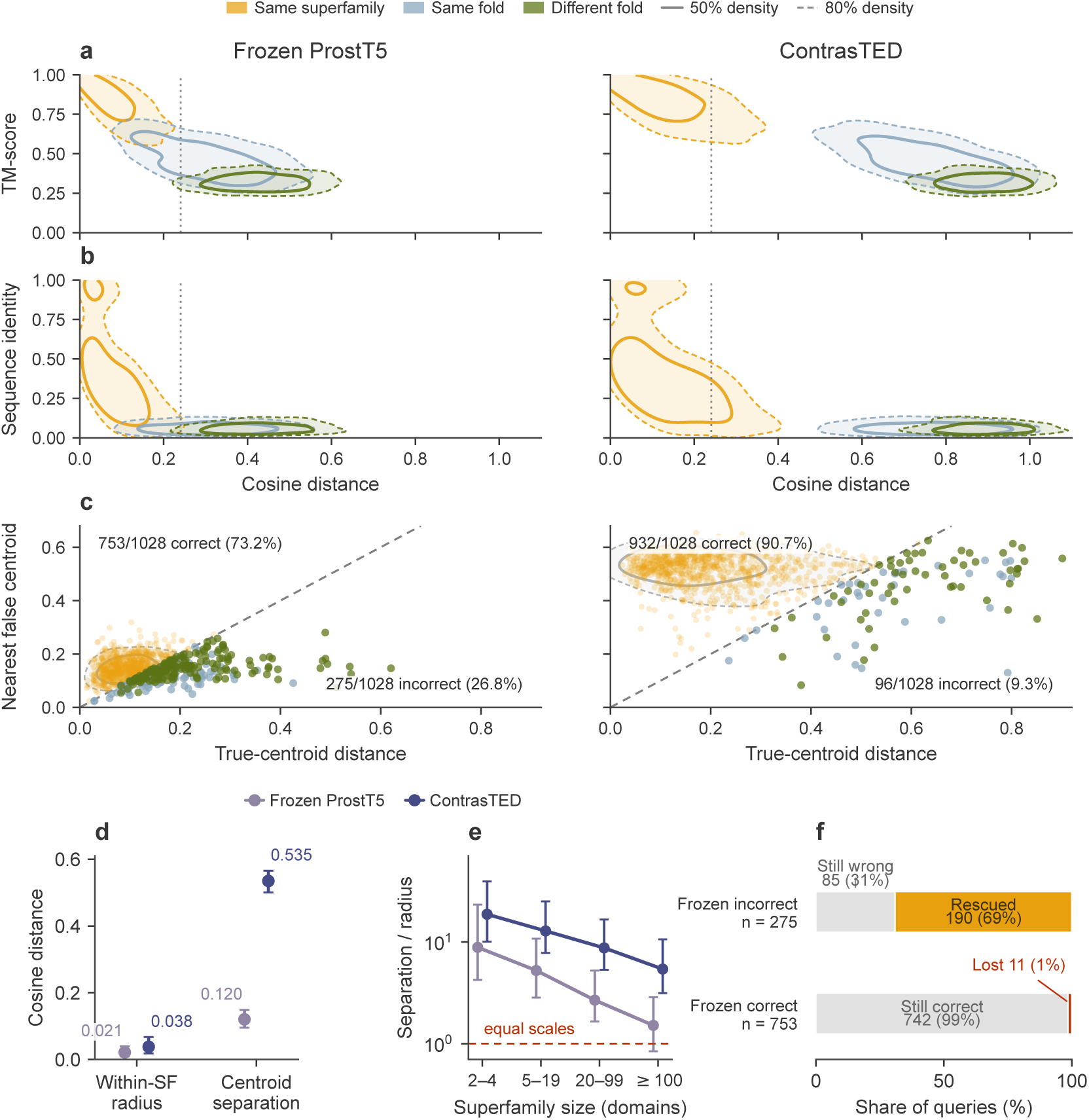
Changes in superfamily margins and remote structural similarity after metric learning. (**a**, **b**) 3,000 balanced domain pairs from 988 topologies. Frozen ProstT5 (left) and ContrasTED (right) cosine distances are plotted against (**a**) TM-score and (**b**) sequence identity. Solid/dashed contours enclose 50%/80% of each class density, and dotted lines mark the calibrated threshold. (**c**) Held-out S20 centroid decisions, plotting labelled-centroid distance (*d*_true_) against nearest-competing-centroid distance (*d*_false_). The diagonal is the exact boundary: correct above, incorrect below; grey contours enclose 50%/80% of query density. Gold, blue and green denote assignment to the true superfamily, a wrong superfamily in the same fold, and a wrong fold, respectively. (**d**) Median within-superfamily radius and nearest-centroid separation across 3,980 training superfamilies with at least two members. (**e**) Median separation/radius across size strata; the red line denotes equal scales. Panels **d** and **e** show interquartile ranges and overlay both representations. (**f**) Paired correctness transitions for all 1,028 queries, normalized within the frozen-incorrect and frozen-correct groups. Gold and red denote changes from incorrect to correct and from correct to incorrect, respectively; grey denotes unchanged correctness.

We next asked whether optimizing for discrete CATH superfamily boundaries distorted the underlying physical relationship between embedding distance and structural similarity. Comparing cosine distance with TM-score across 3,000 balanced domain pairs from 988 topologies revealed that embedding distance remained strongly and monotonically correlated with structural similarity before and after training (*ρ* = *−*0.80 and *−*0.82, paired difference Δ*ρ* = *−*0.020, 95% CI [*−*0.037*, −*0.005]; Fig. 2a). Crucially, this structural correlation was preserved among the 2,197 domain pairs sharing less than 20% sequence identity (*ρ* = *−*0.61; Fig. 2b). ContrasTED therefore sharpens discrete homologous superfamily boundaries while faithfully preserving the continuous structural relationships captured by ProstT5.

Analyzing the 3,980 CATH training superfamilies with at least two members explains these improvements (Fig. 2d,e). Contrastive learning reduced within-superfamily radii (*R*) while increasing nearest-centroid separation (*S*) across all superfamily sizes. This eliminated class overlap (*S/R ≤* 1) and more than doubled the median separation-to-radius ratio across both rare and data-rich superfamilies (Fig. 2d,e).

### 2.3 ContrasTED improves sparsely represented superfamilies

Consistent with the latent-space geometry, the greatest performance advantage occurred among sparsely represented superfamilies (Fig. 3a). For superfamilies in the lowest training-size quartile (1–6 CATH training domains), ContrasTED reached 90.7% H-level accuracy on the S20 test set, compared with 74.9% for frozen ProstT5-AA ∥ 3Di and 79.2% for Foldseek. In contrast, performance across methods converged in the most data-rich quartile (92.5%, 91.8%, and 91.8% for ContrasTED, CATHe2, and Foldseek, respectively). This confirms that proxy-based metric learning effectively resolves the long tail of protein superfamily space, where pairwise alignment and standard classifiers lack sufficient training examples.

**Figure 3:**
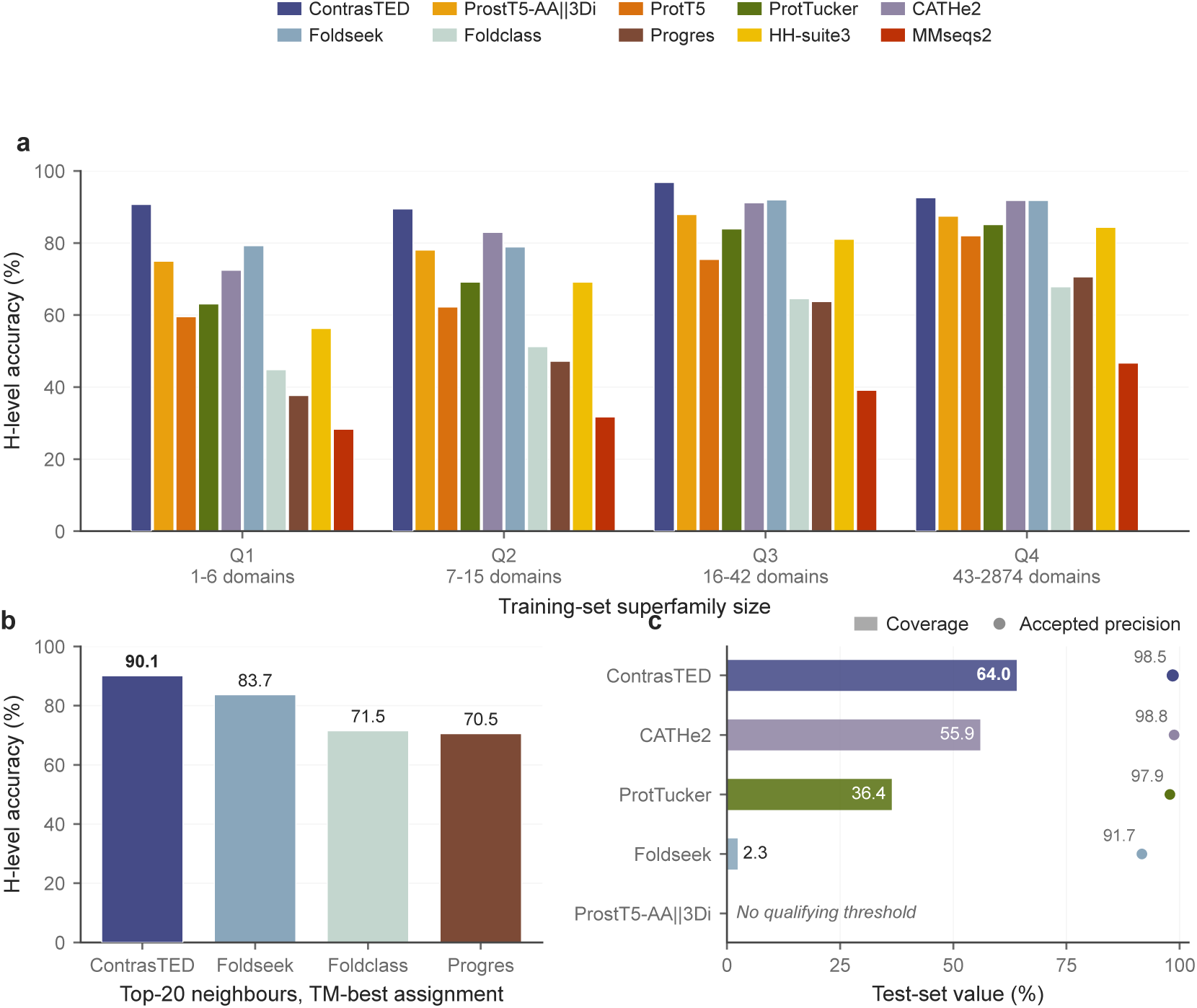
Remote-homology accuracy and confidence-based coverage. (**a**) H-level accuracy by training-set size quartile on the S20 one-per-superfamily remote-homology test set; HH-suite3 searches the train-only TEDLH subset from Table 1. (**b**) H-level accuracy after retrieving the top 20 neighbours and assigning the query-normalized TM-best hit, using the same protocol for ContrasTED, Foldseek, Foldclass, and Progres. (**c**) Test coverage and accepted-set precision after selecting the most permissive threshold with at least 99% empirical precision on validation and applying it to test.

We also compared structure-aware retrieval when each method was allowed 20 neighbours and the hit with the best query-normalized TM-score was taken as the assignment (Fig. 3b). Under this protocol ContrasTED reached 90.1% H-level accuracy, compared with 83.7% for Foldseek, 71.5% for Foldclass, and 70.5% for Progres.

In practical classification pipelines, estimating the reliability of individual predictions is essential for filtering false positives. Because assignment accuracy varied inversely with embedding distance, we calibrated a distance threshold on the validation set by selecting the largest distance retaining at least 99% empirical precision, and applied this cutoff unchanged to the test set (Fig. 3c and Supplementary Table S7). On S20, this validation-calibrated gate retained 64.0% of test queries at 98.5% precision (95% CI [97.2, 99.3]). Under the same 99% validation precision criterion, CATHe2 and ProtTucker retained 55.9% and 36.4% coverage at 98.8% and 97.9% test precision. Foldseek retained 2.3% coverage at 91.7% precision; frozen ProstT5 embeddings had no threshold that reached 99% validation precision. Minor variation between validation and test distributions accounts for the slight difference between the 99% validation selection target and the observed test precision.

Although 1-NN retrieval achieved slightly higher overall accuracy than centroid retrieval on S20 (92.9% vs. 91.4%; Table 1), centroid retrieval performed better when thresholding for high precision. Because 1-NN retrieval relies on the single closest reference domain, occasional boundary outliers cause false positives at moderate distances, requiring a strict threshold (*d ≤* 0.178) that retained only 58.3% test coverage at the 99% precision target (591 correct, 8 errors). In contrast, centroid retrieval averages representations across each superfamily, reducing outlier noise and permitting a more permissive threshold (*d ≤* 0.240) that retained 64.0% coverage (648 correct, 10 errors). Centroid lookup correctly classified 57 additional remote homologues at matched precision while reducing index memory by more than 2,000-fold.

### 2.4 Calibration and database-scale application to TED

Before applying ContrasTED to unannotated domains in TED, we tested whether the distance threshold calibrated on CATH transfers to predicted AlphaFold2 structures. We evaluated the threshold on an independent benchmark of 869 high-confidence domain models with consensus Foldseek and profile-HMM assignments sharing *<* 20% sequence identity to the training set (the TED Gold NR S20 set). Applying the validation-calibrated threshold (*d* = 0.22579) without adjustment retained 688 queries at 98.55% precision (95% CI [97.34, 99.30]; Fig. 4a), confirming that calibration on experimental structures holds on predicted models.

**Figure 4:**
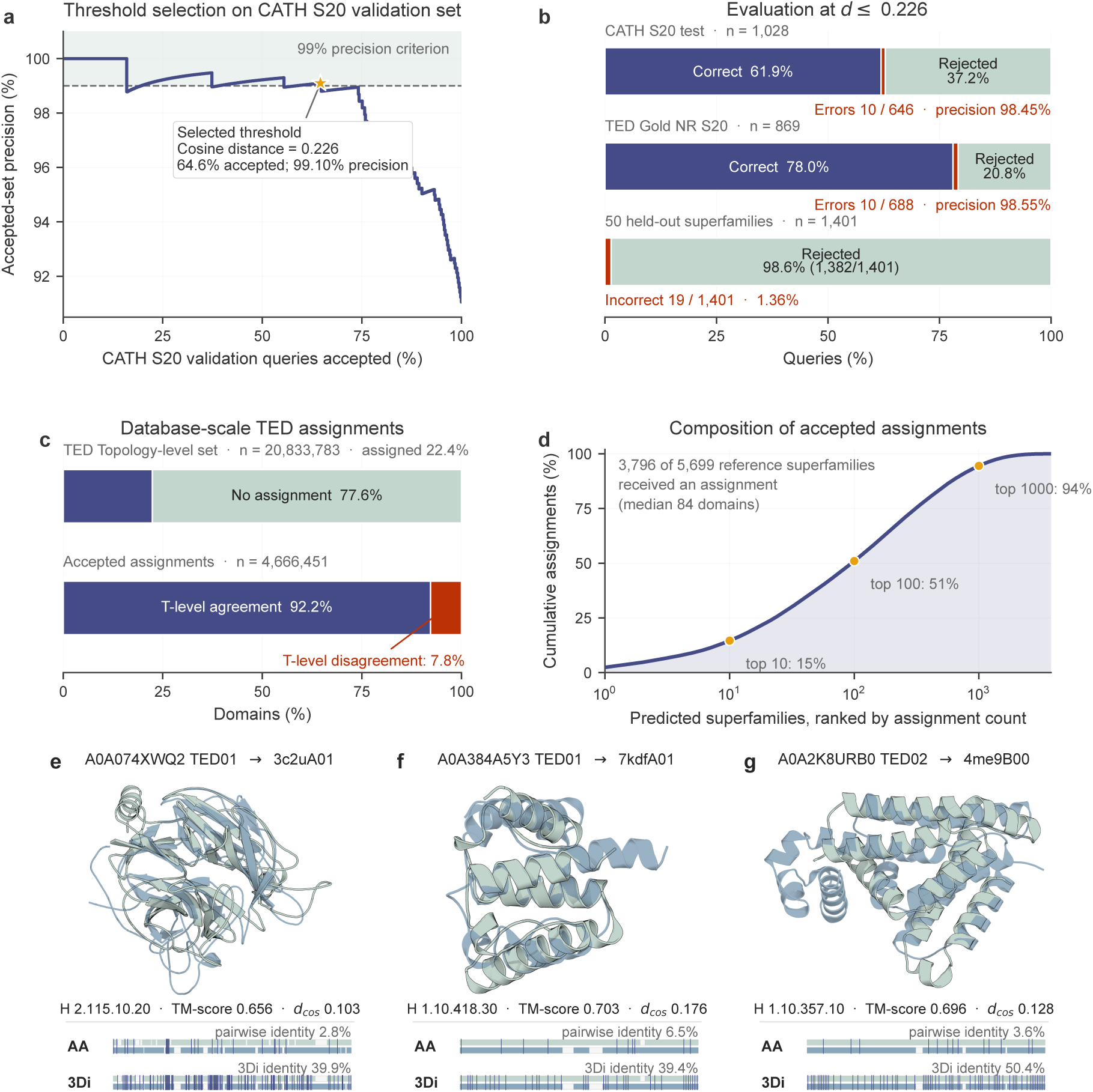
Distance calibration and database-scale application to TED. (**a**, **b**) Separate 50-H S20 model used for absent-superfamily evaluation: its threshold (*d* = 0.22579) was selected at *≥* 99% accepted-set precision on validation and applied to CATH S20 test, TED Gold NR S20 and 50 superfamilies withheld from training and lookup. (**c**–**g**) ContrasTED-AA ∥ 3Di model, using its calibrated precision threshold (*d* = 0.24048). (**c**) Assignment yield and T-level consistency for 20,833,783 TED domains with Topology assignments. (**d**) Cumulative assignment share by ranked superfamily. (**e**) The AF-A0A074XWQ2-F1-model v4 TED01 assignment discussed in the text. (**f**, **g**) Two further remote assignments with TM-align support. All three pass the calibrated precision threshold (*d* = 0.240), have TM-align support, 2.8–6.5% sequence identity and no H-level assignment from the Foldseek-based TED procedure. Mint and sky blue denote TED and CATH alignment coverage; navy ticks denote identical residues or 3Di states. 3Di identity is the same-state paired fraction; TM-score uses the shorter domain.

We also tested whether the threshold reliably rejects structures from superfamilies not present in CATH. We trained a separate model after withholding 50 CATH superfamilies from both training and reference sets (Methods). Applying its validation-calibrated threshold (*d* = 0.22579) correctly rejected 1,382 of 1,401 queries from the withheld superfamilies (98.64%; Fig. 4b), with false acceptances concentrated in just two groups (Supplementary Table S10). Distance thresholding therefore reliably rejects unseen superfamilies while maintaining high sensitivity to divergent homologues.

Having verified threshold performance on predicted structures and unseen superfamilies, we computed reference centroids across all 5,659 curated CATH superfamilies in classes 1–3 and applied ContrasTED to 20,833,783 TED domains that had received a CATH Topology label from structural search but lacked an H-level assignment. ContrasTED assigned 4,666,451 domains (22.40%) to 3,796 homologous superfamilies with high confidence (Fig. 4c). Crucially, 92.22% of these predicted H assignments (4,303,506 domains) agreed with the existing TED Topology, providing independent hierarchical support. Assignments spanned both common and rare superfamilies, with 696 superfamilies receiving at least 1,000 new domain members (Fig. 4d). Database-scale classification is fast once embeddings are generated. While paired ProstT5 embedding generation required 9.8–10.4 ms per domain on a single NVIDIA L40S GPU (*∼*100 domains/s per GPU, parallelizing linearly across multi-GPU nodes), downstream projection and nearest-centroid search against all 5,659 CATH superfamilies exceeded 100,000 queries/s on a single CPU core against a 1.4 MB index (Supplementary Table S4).

Individual assignments illustrate how ContrasTED recovers remote evolutionary relationships deep in the sequence twilight zone. For example, ContrasTED assigned the uncharacterized domain AF-A0A074XWQ2-F1-model v4 TED01 to CATH superfamily 2.115.10.20 at a cosine distance of 0.103 (well within the *d* = 0.240 threshold), matching the CATH representative 3c2uA01 (Fig. 4e). Pairwise sequence identity between these domains was only 2.8%—far below the detection limit of sequence alignment methods—and previous structural search tools used in TED were unable to detect this relationship or make an H-level assignment. Despite this extreme sequence divergence, structural superposition with TM-align revealed clear structural homology, aligning 251 residues with a TM-score of 0.656 and an RMSD of 3.3 Å (Fig. 4e). Additional examples with sequence identities between 2.8% and 6.5% showed similarly strong structural alignment (TM-scores *>* 0.6; Fig. 4f,g). Panel 4g assigns AF-A0A2K8URB0-F1-model v4 TED02 to 1.10.357.10 at *d* = 0.128 (TM-score 0.696, 125 residues, RMSD 2.74 ^°^A, 3.6% identity).

## 3 Discussion

Organizing predicted protein structures into evolutionary superfamilies requires methods that remain sensitive at extreme sequence divergence while scaling to millions of domains. While contrastive learning can adapt protein language models to structural hierarchies, pairwise and triplet-based objectives struggle with the long tail of rare superfamilies, where positive pairs rarely co-occur during training. ContrasTED addresses this by combining multimodal ProstT5 representations (AA ∥ 3Di) with center-contrastive learning, anchoring each superfamily to a learnable proxy center. This yields both high classification accuracy on rare superfamilies and rapid centroid-based search across whole databases, complementing residue-level alignment tools such as profile HMMs and EBA that provide detailed evolutionary alignments for site-level interpretation [24, 27].

The performance gains reflect three complementary factors: multimodal input, metric learning, and dataset scale. First, adding 3Di structural tokens to amino-acid embeddings added 3.4 percentage points to frozen retrieval, bringing it near parity with Foldseek. Second, center-contrastive training on CATH and TED domains brought overall H-level accuracy to 92.9%. Local structural tokens provide backbone context, metric learning separates superfamily boundaries, and TED augmentation supplies the broad sequence diversity needed for remote homology.

The greatest accuracy gains occurred in rare superfamilies with only one or two CATH training domains, where standard classifiers and structure search methods lose sensitivity. While triplet-based methods require multiple members of a superfamily to co-occur in the same batch to form positive pairs, ContrasTED compares every domain against the full bank of learnable centers. Combined with balanced superfamily sampling, this ensures that singleton and rare superfamilies generate direct gradient signals in every update rather than being overshadowed by large families.

Embedding distance correlates strongly with TM-score (*ρ* = *−*0.82 overall and *ρ* = *−*0.61 below 20% sequence identity; Fig. 2a,b), showing that the model preserves physical structural similarity. High TM-scores remain a trusted indicator of homology, providing independent confirmation for remote predictions in TED (Fig. 4e–g). However, supervised metric learning provides greater sensitivity than structural search tools. Even when Foldseek selected the candidate with the highest TM-score among 20 hits, ContrasTED achieved higher classification accuracy (90.1% vs. 83.7%; Fig. 3b). This allows ContrasTED to reflect continuous structural similarity while clearly separating distinct evolutionary superfamilies.

Embedding distance serves as a reliable measure of prediction confidence. Calibrating a threshold on validation data allowed high-precision classification while rejecting unrepresented superfamilies. Centroid retrieval proved particularly advantageous under this high-confidence regime. While 1-NN search relies on single nearest neighbours that can be sensitive to isolated reference outliers, superfamily centroids average across member domains, smoothing local noise and yielding higher coverage under calibrated precision targets. Applying this threshold to 20.8 million unassigned TED domains generated 4.67 million high-confidence superfamily assignments across 3,796 superfamilies, with 92.2% topological agreement with existing annotations.

Several methodological aspects should be considered. First, training incorporates consensus labels transferred from TED, which could carry residual noise from automated pipelines. Second, ContrasTED relies on predefined domain boundaries and focuses on CATH classes 1–3. Finally, while nearest-centroid lookup provides rapid global classification, it does not generate residue-level alignments; hence, ContrasTED and alignment-based tools serve complementary roles in structural bioinformatics.

ContrasTED combines multimodal protein embeddings with center-contrastive learning to deliver an accurate, scalable framework for remote-homology classification across CATH. By eliminating the batch-sampling constraints of pairwise objectives on rare superfamilies and enabling rapid centroid search, ContrasTED achieves higher sensitivity than fast structural search tools while maintaining the throughput needed to organize millions of predicted protein structures.

## 4 Methods

### 4.1 Base embeddings from protein language models

We used ProstT5 [10] as the primary feature extractor. For each protein domain, final-layer hidden states were mean-pooled across residue positions under both the amino-acid mode (`<AA2fold>`) and the 3Di structural mode (`<fold2AA>`), excluding special tokens. Each mode produces a 1,024-dimensional embedding; concatenating the two vectors yields the 2,048-dimensional AA ∥ 3Di domain representation [29]. We extracted 3Di token sequences with Foldseek from domain PDB structures for TED entries and from experimental coordinates for CATH benchmark domains. Domains shorter than 16 residues or longer than 2,000 residues were excluded; nonstandard amino acids were mapped to X; and discontinuous CATH domains with multiple segments were concatenated in chain order. Single-modality ProstT5 and ProtT5 embeddings [6] were extracted under identical pooling protocols as benchmark controls (Table 1).

### 4.2 ContrasTED encoder architecture

The trainable ContrasTED projection head *g_ϕ_* is a two-layer multilayer perceptron (MLP). The input layer maps 2,048-dimensional AA ∥ 3Di embeddings to a 512-dimensional hidden representation with 1D batch normalization, GELU activation [11], and dropout of 0.1 (using accumulated running statistics at inference). The second linear layer projects the hidden representation to 128 dimensions, followed by L2 normalization (∥ **z** ∥ _2_ = 1). Cosine distance *d*(**u**, **v**) = 1 *−* **u**^T^**v** is therefore the operative metric for retrieval and classification. Reported ContrasTED-AA ∥ 3Di benchmark results in Table 1 are the mean of seeds 40–42. Single-seed analyses and figures use the canonical seed-40 checkpoint.

### 4.3 Center-contrastive learning

We used center-contrastive loss (CCL) as a proxy-based metric learning objective [3]. Let a mini-batch contain *N* L2-normalized domain embeddings **z***_i_ ∈* ℝ^128^ with CATH homologous superfamily labels *y_i_ ∈ {*1*, … , K}*. The objective maintains a bank of *K* learnable parameter vectors, where each normalized vector **c***_j_* (∥ **c***_j_* ∥ _2_ = 1) acts as a proxy center for superfamily *j*. For each domain–center pair (*i, j*), the scaled and margin-penalized similarity *u_ij_* and its softmax probability *p_ij_* are defined as:

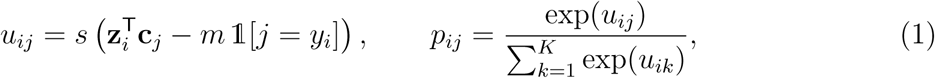

where *s* is a scaling factor and *m* is an additive cosine margin applied exclusively to the true class center. Incorporating label smoothing *ε*, the target distribution *q_ij_* assigns 1 *− ε* to the ground-truth class (*j* = *y_i_*) and *ε/*(*K −* 1) uniformly across all competing centers. The total batch loss combines cross-entropy over proxy similarities with an L2 center regularization penalty:

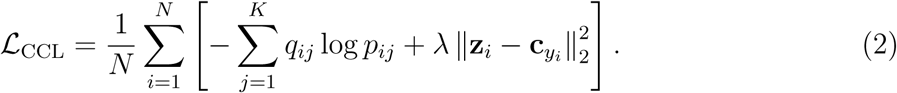

We trained all models with *s* = 32, *m* = 0.3, *λ* = 1.0, and *ε* = 0.1.

By evaluating an *N × K* similarity matrix against all superfamily proxies simultaneously, every sampled domain generates informative gradient updates relative to the full superfamily roster. Mini-batches were constructed by sampling superfamilies uniformly without replacement and drawing at most two domains per superfamily, yielding 1,024 domains per batch (112 batches per epoch on the full training set). Singleton superfamilies (represented by only one CATH training domain) contribute direct gradient updates under this formulation. Crucially, the learnable proxy centers **c***_j_* were used only during optimization; at inference, retrieval centroids were computed as empirical L2-normalized means of projected CATH reference embeddings.

### 4.4 Datasets and benchmark construction

Training, validation, and test splits were derived from CATH v4.4 classes 1–3 (Mainly Alpha, Mainly Beta, and Alpha-Beta). For the S20 benchmark, we sampled one representative domain per superfamily into disjoint validation and test sets. To enforce strict sequence separation, CATH S100 domains sharing *≥* 20% sequence identity to any validation or test domain were removed using MMseqs2 profile search (sensitivity 7.5, three iterations, and 80% bidirectional coverage) [28]. Test superfamilies with no remaining training representatives after filtering were excluded. The resulting S20 benchmark comprises 116,301 training, 514 validation, and 1,028 test domains across 5,659 training superfamilies. In all Table 1 evaluations, the reference lookup set contains only the corresponding CATH training domains.

To expand the training set with structural diversity from AlphaFold DB, we augmented CATH supervision with domain representatives from The Encyclopedia of Domains (TED) [16]. Within TED, high-confidence domain annotations are defined by consensus assignments supported by both Foldseek structural search and profile-HMM search (the “TED Gold” set). We extracted S40 representatives from this high-confidence set and applied MMseqs2 sequence profile filtering to remove any domain sharing *≥* 20% sequence identity to the CATH S20 validation or test sets, yielding 11,131,930 filtered TED training domains. These domains were used exclusively for metric-learning data augmentation; the lookup set remained strictly CATH-only, ensuring that benchmark evaluations measure annotation transfer from curated experimental structures. An independent, non-redundant S20 subset of TED Gold domains sharing *<* 20% sequence identity to the entire combined CATH+TED training set was withheld from training and reserved for external validation.

### 4.5 Training and optimization

We trained the S20 model on the combined CATH+TED dataset, selecting the final model checkpoint by 1-NN top-1 validation accuracy against the CATH training set. Amino-acid-only baseline models followed the same training and checkpoint selection protocol. Full optimization hyperparameters are provided in the Supplementary Methods.

### 4.6 Embedding-space analyses

Embedding geometry analyses (Fig. 2) used the S20 AA ∥ 3Di model and 3,000 balanced domain pairs sampled across 988 CATH topologies. Pairs were drawn from CATH S95 representatives in three balanced strata: (i) same homologous superfamily (*n* = 1,000), (ii) same Topology but distinct superfamily (*n* = 1,000), and (iii) different Topology (*n* = 1,000). Structural alignment scores were computed with TM-align [33, 34], and pairwise sequence identities were computed by Smith–Waterman local alignment (BLOSUM62 matrix, affine gap penalties of *−*11 open and *−*1 extend), each normalized by the shorter domain length. Spearman rank correlation confidence intervals were estimated by block-bootstrapping over complete topologies. For each training superfamily with at least two members, within-class radius *R* was calculated as the median cosine distance of member domains to the class centroid, and separation *S* as the cosine distance to the nearest competing centroid. For held-out test queries, the centroid decision margin was defined as *d*_false_ *− d*_true_, and paired margin changes before and after contrastive learning were evaluated using 2,000 bootstrap resamples.

### 4.7 Comparative evaluation

Top-1 accuracy was computed as the fraction of test queries assigned to the correct homologous superfamily, treating unanswered queries as incorrect. Because each test set contains exactly one query per superfamily, top-1 accuracy is directly equivalent to macro recall across superfamilies. We evaluated sensitivity to remote homology under a shared structural-ranking protocol by retrieving 20 neighbours per method and assigning the hit with the highest query-normalized TM-score.

Evaluated baseline methods comprise: (i) sequence search via MMseqs2 [28]; (ii) profile HMM search with HH-suite3 against a TEDLH subset [2, 27]; (iii) frozen embeddings from ProtT5 and ProstT5 (1-NN and nearest-centroid) [6, 10]; (iv) the CATHe2 supervised neural network classifier [20]; (v) the ProtTucker contrastive baseline [9]; and (vi) structure retrieval tools including Foldseek, Foldclass, and Progres [7, 15, 29]. CATHe2 was retrained on the identical CATH splits and multimodal inputs. ProtTucker was evaluated as an external baseline using its published CATH v4.3 checkpoint. For Table 1 we searched a TEDLH subset containing only profiles whose CATH seed domain belongs to the training split. This prevents a query from matching a profile seeded on itself and removes profiles seeded on domains withheld from training by the S20 identity filter, matching the CATH-train lookup used by the other methods. Full comparator parameters, reachability, and library variants are detailed in Supplementary Tables S6 and S8–S9.

### 4.8 Annotation transfer

For Table 1, we indexed projected training-set embeddings and evaluated both 1-NN retrieval and nearest-centroid retrieval (using the L2-normalized means of projected CATH training domains for each superfamily). For confidence-gated evaluation (Fig. 3c), each method was ranked by its native confidence metric: cosine distance for ContrasTED and frozen embeddings, maximum softmax probability for CATHe2, and E-value for Foldseek. For each method, we selected the most permissive validation threshold achieving *≥* 99% precision on the S20 validation set and applied that cutoff unchanged to the test set. Exact Clopper–Pearson binomial confidence intervals were computed on accepted test predictions.

### 4.9 Uncertainty and statistical analysis

Benchmark accuracy confidence intervals were computed using 2,000 percentile bootstrap resamples of the test set. Because each remote-homology test set contains one query per superfamily, resampling queries is equivalent to resampling superfamilies. Differences in accuracy and held-out centroid margins were evaluated with paired query bootstraps; Spearman correlation intervals resampled complete Topology blocks; and precision and rejection intervals were calculated using exact Clopper–Pearson binomial limits. All model hyperparameters and distance thresholds were selected exclusively on training and validation sets before evaluating test data.

### 4.10 Runtime and database-scale evaluation

ProstT5 fp16 inference latency was measured on an NVIDIA L40S GPU (excluding initial model loading), with separate passes for amino-acid and 3Di modes and including Foldseek 3Di extraction time. The 2,048-to-128 projection head was benchmarked across 119,113 domains on Apple Silicon (MPS). Nearest-centroid search was benchmarked using 10,000 queries against all 5,659 CATH superfamily centroids using FAISS (float32 search) on a single Apple arm64 CPU core [4]; index size reports the centroid matrix in fp16 precision (Supplementary Table S4).

#### Annotation of unassigned TED domains

The unassigned TED dataset comprises 20,833,783 domains that received a CATH Topology assignment from the TED structural search pipeline but lacked an H-level superfamily assignment. This set includes 16,026,530 Foldseek fold-level hits and 4.8 million remainder hits identified by Merizo-search and Foldclass (TM-align *>* 0.5 normalized by domain length) [16]. For database-scale annotation, we used the ContrasTED-AA ∥ 3Di model with nearest-centroid retrieval against all 5,659 homologous-superfamily centroids across CATH classes 1–3. The distance threshold (*d* = 0.24048) was calibrated on the 514 CATH S20 validation queries to achieve *≥* 99% empirical precision, and fixed before scoring the 1,028 CATH S20 test queries and the unassigned TED dataset.

#### Out-of-distribution and TED Gold evaluation

To evaluate out-of-distribution rejection, we trained a separate S20 model after removing 50 CATH superfamilies from the CATH and TED training sets and from the reference lookup set. Withheld superfamilies yielded 1,401 absent-class domains. The distance threshold (*d* = 0.22579) was calibrated on the standard validation set to achieve *≥* 99% precision and applied to the withheld superfamilies to measure the proportion correctly rejected beyond the cutoff (Supplementary Table S10). For external validation on divergent known superfamilies, we used the TED Gold NR S20 benchmark. Starting from TED Gold domains (which share consensus Foldseek and profile-HMM assignments), we removed domains sharing *≥* 20% sequence identity to the complete CATH+TED training set. Retaining one representative domain per superfamily yielded 869 sequence-remote test queries, which were evaluated under the validation-calibrated distance threshold without further adjustment.

## Acknowledgements

We thank the members of the Orengo group who provided feedback on the benchmark design, manuscript framing, and interpretation of the results.

## Author contributions

D.M. conceptualized and designed the ContrasTED method, CATH benchmark and TED annotation analyses. D.M. implemented the training and inference software. D.M. carried out all model training and comparative benchmark experiments. D.M. produced the figures and tables. D.M. wrote the manuscript with input from all coauthors. M.H., N.B. and C.O. contributed to conceptualization of the study and to writing the manuscript. V.W. contributed to structural analysis and data curation. J.J. contributed to data analysis.

## Data availability

The public resources analyzed in this study are available from the original CATH and TED releases cited in the manuscript. Derived datasets supporting the findings of this study, including the S20 benchmark split, benchmark model predictions, checkpoint hashes, figure source data and TED T*→*H assignment summaries, are assembled in a versioned archive for release with the preprint.

## Code availability

Source code for ContrasTED is publicly available without access restrictions at https://github.com/dmmiller597/contrasted. The repository contains the training and inference package. A tagged release and corresponding versioned archive matching this manuscript revision will be linked from the repository when the preprint is posted and its DOI is minted.

