## supplementary for "ContrasTED: contrastive domain embeddings for scalable remote homology classification"

Supplementary Information

#### Supplementary Tables

Table S1: **Full CATH-level accuracies for main-text Table 1.** Each cell stacks accuracy (%) above a 95% bootstrap percentile interval (2,000 resamples over test queries). Accuracy uses the full S20 test roster ($n$=1028; unanswered Foldseek queries count as errors). A blank method name repeats the method above under centroid retrieval. For the three-seed ContrasTED-AA‖3Di rows, correctness is first averaged across the three predictions for each query and that query-level mean is bootstrapped; other rows bootstrap the single reported checkpoint. Methods and footnotes follow main-text Table 1.

| Method | Mode | C | A | T | H |
|---|---|---|---|---|---|
| *S20 split ($n = 1028$)* | | | | | |
| *Sequence* | | | | | |
| MMseqs2 | 1-NN | 39.5 [36.6–42.5] | 37.9 [35.0–41.0] | 37.3 [34.3–40.3] | 36.3 [33.3–39.2] |
| HH-suite3† | profile | 85.7 [83.5–87.7] | 78.6 [76.1–81.0] | 74.7 [72.1–77.4] | 72.3 [69.7–74.9] |
| *Structure* | | | | | |
| Foldseek (default)‡ | 1-NN | 94.5 [93.1–95.8] | 92.0 [90.4–93.7] | 89.3 [87.4–91.1] | 85.3 [83.2–87.5] |
| Foldseek (sensitive) | 1-NN | 96.5 [95.3–97.6] | 94.5 [93.0–95.8] | 92.5 [90.9–94.1] | 88.2 [86.2–90.3] |
| Progres | 1-NN | 95.7 [94.5–96.9] | 81.2 [79.0–83.7] | 69.5 [66.8–72.5] | 54.4 [51.5–57.4] |
| Foldclass | 1-NN | 96.5 [95.3–97.5] | 85.1 [83.1–87.2] | 72.8 [70.2–75.4] | 56.8 [53.8–59.9] |
| *Frozen embeddings* | | | | | |
| ProtT5 | 1-NN | 93.0 [91.4–94.6] | 83.9 [81.4–86.1] | 75.5 [72.8–78.1] | 69.6 [66.8–72.3] |
| ProstT5-AA | 1-NN | 96.3 [95.1–97.4] | 90.3 [88.4–92.0] | 85.3 [83.1–87.5] | 78.5 [76.1–81.0] |
| | centroid | 94.8 [93.5–96.1] | 85.2 [83.1–87.3] | 77.3 [74.9–79.9] | 66.8 [63.9–69.6] |

*Continued on next page*

Table S1: **Full CATH-level accuracies for main-text Table 1** (continued).

| Method | Mode | C | A | T | H |
|---|---|---|---|---|---|
| ProstT5-AA‖3Di | 1-NN | 97.6 [96.5–98.4] | 92.1 [90.4–93.7] | 88.4 [86.4–90.4] | 81.9 [79.6–84.3] |
| | centroid | 96.0 [94.7–97.2] | 89.7 [87.8–91.4] | 83.9 [81.6–86.2] | 73.2 [70.6–76.0] |
| *Supervised ANN* | | | | | |
| CATHe2-AA§ | softmax | 96.5 [95.3–97.6] | 93.0 [91.4–94.6] | 87.6 [85.7–89.7] | 82.1 [80.0–84.5] |
| CATHe2-AA‖3Di§ | softmax | 98.2 [97.4–99.0] | 95.3 [94.0–96.5] | 91.7 [90.0–93.4] | 84.2 [82.1–86.5] |
| *Contrastively projected* | | | | | |
| ProtTucker | 1-NN | 95.1 [93.8–96.4] | 88.2 [86.2–90.2] | 82.0 [79.7–84.4] | 75.0 [72.4–77.6] |
| ContrasTED-AA | 1-NN | 97.2 [96.1–98.2] | 93.5 [91.9–94.9] | 91.1 [89.4–92.6] | 88.7 [86.9–90.7] |
| | centroid | 97.0 [95.9–98.0] | 92.8 [91.1–94.4] | 90.2 [88.4–91.8] | 87.5 [85.4–89.5] |
| ContrasTED-AA‖3Di (mean) | 1-NN | 98.9 [98.4–99.4] | 97.3 [96.4–98.1] | 95.9 [94.7–96.9] | 92.9 [91.4–94.3] |
| | centroid | 98.3 [97.6–98.9] | 96.5 [95.5–97.4] | 95.1 [93.9–96.2] | 91.4 [89.9–93.0] |

Table S2: **Seed stability of production S20 CATH+TED ContrasTED-AA‖3Di.** Test homologous-superfamily (H) accuracy (%) on the remote-homology S20 split ($n$=1028) under 1-NN and nearest-centroid retrieval. Training mixes CATH with sequence-filtered TED domains; lookup is the CATH training set only (same protocol as main-text Table 1). Seeds 40, 41 and 42 share hyperparameters (lr 0.012, scale 32, margin 0.3, $\lambda$=1.0, early stopping on validation 1-NN against the CATH training set). Main-text Table 1 reports the three-seed mean; main-text Figures 2–4 use seed 40. Mean ± s.d. over the three seeds.

| Modality | Mode | Seed 40 | Seed 41 | Seed 42 | Mean ± s.d. |
|---|---|---|---|---|---|
| AA‖3Di | 1-NN | 92.3 | 93.4 | 92.9 | 92.9 ± 0.5 |
| | centroid | 90.7 | 91.5 | 92.1 | 91.4 ± 0.7 |

Table S3: **Contribution of contrastive projection and TED training augmentation.** Homologous-superfamily (H) accuracy (%) on the remote-homology S20 split ($n$=1028) under 1-NN and nearest-centroid retrieval. Lookup is the CATH training set only (same protocol as main-text Table 1). Rows compare raw ProstT5-AA‖3Di embeddings, a center-contrastive head trained on CATH alone, and the production head trained on the CATH+TED mixture. TED domains enlarge the training mixture but never enter the lookup set. All trained rows use seed 40, so the CATH+TED cells are the seed-40 components of the three-seed means in main-text Table 1.

| Encoder | 1-NN | Centroid |
|---|---|---|
| Raw ProstT5-AA‖3Di | 81.9 | 73.2 |
| CCL (CATH train) | 87.8 | 86.4 |
| CCL (CATH+TED train) | 92.3 | 90.7 |

Table S4: **Production annotation cost.** ProstT5 wall-clock time was measured on an NVIDIA L40S with fp16 inference; model loading is excluded. AA and 3Di require separate encoder passes. Foldseek 3Di extraction contributes less than 0.1 ms/domain and is included. The production 2048-to-128 projection was measured on Apple MPS, and float32 centroid lookup used FAISS on an Apple arm64 CPU [2]; index size reports fp16 storage. Projection and lookup use a prebuilt CATH reference; one-time database construction is excluded.

| Stage / input | Passes or index | Throughput / cost |
|---|---|---|
| ProstT5 AA (`<AA2fold>`) | 1 pass | 4.9–5.2 ms/domain |
| ProstT5 3Di (`<fold2AA>`) | 1 pass | 4.9–5.2 ms/domain |
| ProstT5 AA‖3Di | 2 passes | 9.8–10.4 ms/domain |
| Projection head | 2048→128 | ~98,000 domains/s |
| Nearest-centroid lookup | 5,659 H classes | >100,000 queries/s |
| Prebuilt centroid index | 128-d, fp16 | 1.4 MB |

Table S5: **Locked production training configuration.** Values are those of the S20 AA‖3Di production encoder unless stated otherwise. The runtime number of classes is the number of H labels in the training mixture (5,659); the loss uses a dense learnable center bank of that size.

| Component | Setting |
|---|---|
| Input / projection | 2048 → 512 → 128; BN, GELU, dropout 0.1; L2-normalized output |
| Objective | Center-contrastive at CATH H level |
| Loss parameters | $s = 32$, $m = 0.3$, $\lambda = 1.0$, $\varepsilon = 0.1$ |
| Batch construction | 1024 examples; class-balanced; at most 2 per H class; 112 batches/epoch |
| Optimizer | AdamW [4]; lr 0.012; weight decay $10^{-4}$; gradient clip 1.0 |
| Schedule | 10-epoch linear warm-up; cosine decay to $10^{-6}$; maximum 80 epochs |
| Selection | validation 1-NN H accuracy against the CATH training set; patience 15 |
| Numeric mode / seed | bf16 mixed precision; deterministic; seed 40 |

Table S6: **Structure-search H-level accuracies** on the same remote-homology S20 query set as main-text Table 1 ($n$=1028). Headline Table 1 uses Foldseek in default mode ($-s\,9.5$, `--max-seqs` 1,000, top hit by bitscore) and sensitive mode (`--exhaustive-search 1`), and Foldclass and Progres with 1-NN retrieval. ContrasTED rows reproduce the three-seed AA‖3Di means from Table 1. Accuracies are overall H-level; unanswered queries count as errors.

| Method | Setting | H S20 (%) |
|---|---|---|
| Foldseek (default) | $-s\,9.5$, `--max-seqs` 1,000 | 85.3 |
| Foldseek (sensitive) | `--exhaustive-search 1` | 88.2 |
| Progres (Table 1) | 1-NN | 54.4 |
| Foldclass (Table 1) | 1-NN | 56.8 |
| ContrasTED-AA‖3Di (Table 1) | 1-NN | 92.9 |
| ContrasTED-AA‖3Di (Table 1) | centroid | 91.4 |

Table S7: **Checkpoint-specific thresholds selected at 99% validation precision.** Each row uses nearest-centroid retrieval and the maximum observed validation distance retaining at least 99% point precision among accepted assignments. Numerical thresholds are selected independently because distance scales differ between trained models. The full S20 row is the threshold used for TED T-hit assignment in main-text Figure 4c–g. The 50-H row uses the same rule on a separately trained encoder; the held-out superfamilies were excluded from training and lookup and did not enter threshold selection.

| Model | Split | $d$ | Val. prec. (%) | Val. cov. (%) | Test prec. (%) | Test cov. (%) |
|---|---|---|---|---|---|---|
| Full CATH+TED, seed 40 | S20 | 0.24048 | 99.14 | 68.09 | 98.48 | 64.01 |
| 50-H holdout, seed 40 | S20 | 0.22579 | 99.10 | 64.59 | 98.45 | 62.84 |

Table S8: **Matched comparison on the TEDLH-reachable subset.** All Table 1 methods restricted to the S20 test queries whose superfamily is present in the train-only TEDLH profile subset searched by HH-suite3 ($n = 959$ of 1,028). Every query in the subset enters the denominator (unanswered queries count as errors).

| Method | Mode | Accuracy (%; S20) | | | |
|---|---|---|---|---|---|
| | | C | A | T | H |
| *Sequence search* | | | | | |
| MMseqs2 | 1-NN | 39.6 | 38.0 | 37.2 | 36.4 |
| HH-suite3[¶] | profile | 89.1 | 82.8 | 79.6 | 77.5 |
| *Structure search* | | | | | |
| Foldseek (default)[‡] | 1-NN | 96.0 | 93.5 | 90.8 | 87.1 |
| Foldseek (sensitive) | 1-NN | 97.2 | 95.2 | 93.1 | 89.2 |
| Progres | 1-NN | 95.8 | 81.5 | 69.4 | 55.6 |
| Foldclass | 1-NN | 96.5 | 85.0 | 72.4 | 57.8 |
| *Frozen pLM embeddings* | | | | | |
| ProtT5 | 1-NN | 93.5 | 84.7 | 77.0 | 71.0 |
| ProstT5-AA | 1-NN | 96.9 | 91.2 | 86.8 | 80.3 |
| | centroid | 95.2 | 86.2 | 78.4 | 68.3 |
| ProstT5-AA‖3Di | 1-NN | 97.5 | 92.5 | 88.8 | 83.0 |
| | centroid | 95.9 | 89.7 | 83.9 | 73.9 |
| *Supervised ANN* | | | | | |
| CATHe2-AA[§] | softmax | 96.7 | 93.6 | 89.1 | 83.8 |
| CATHe2-AA‖3Di[§] | softmax | 98.4 | 95.6 | 92.3 | 85.5 |
| *Contrastively projected embeddings* | | | | | |
| ProtTucker | 1-NN | 95.6 | 89.6 | 83.8 | 77.2 |
| ContrasTED-AA | 1-NN | 97.7 | 94.5 | 92.6 | 90.7 |
| | centroid | 97.4 | 93.8 | 91.6 | 89.3 |
| ContrasTED-AA‖3Di | 1-NN | 99.0 | 97.6 | 96.0 | 93.6 |
| | centroid | 98.0 | 96.5 | 94.8 | 91.9 |

Table S9: **HH-suite3 searches of TEDLH library variants.** The same HH-suite3 searches scored against progressively stricter TEDLH libraries, on the full S20 test roster ($n$=1028). TEDLH profiles are keyed by the CATH domain on which they were seeded, so library membership can be checked directly. Lite and full libraries contain test-domain seeds, including the query's own profile, and are not comparable to Table 1. Self-exclusion removes only that self-hit; profiles seeded on other domains withheld from training by the S20 identity filter remain. Only the train-only row matches the CATH-train lookup used throughout Table 1; it is the row reported there.

| Library | Answered | Accuracy (%; S20) | | | |
|---|---|---|---|---|---|
| | | C | A | T | H |
| TEDLH lite<br>*50k profiles; contains test domains* | 926 | 83.6 | 78.1 | 75.7 | 74.1 |
| TEDLH full<br>*934k profiles; contains test domains* | 1025 | 93.4 | 90.8 | 89.3 | 88.6 |
| TEDLH full, self-excluded<br>*query's own profile removed* | 1024 | 88.0 | 82.2 | 78.7 | 76.8 |
| TEDLH train-only<br>*training-split seeds only* | 1024 | 85.7 | 78.6 | 74.7 | 72.3 |

Table S10: **Per-superfamily rejection in the frozen 50-H absent-reference cohort.** The 50 CATH H groups were sampled from train-only superfamilies containing 11–99 domains (seed 42) and then fixed for the S20 experiment. The seed-40 H50 model removed these groups from both CATH and TED training and from lookup. The distance threshold ($d$=0.22579) was selected on the standard S20 validation set without using this cohort. Overall, 1,382/1,401 domains were rejected (98.64% micro-average; 97.07% macro-average across H groups), and 48/50 superfamilies had no false accept. The 19 false accepts are confined to 2.30.31.40 (12/12) and 3.10.450.360 (7/15).

| CATH H | Domains | False accepts | Rejected (%) | Median $d$ |
|---|---|---|---|---|
| 1.10.100.10 | 15 | 0 | 100.0 | 0.452 |
| 1.10.1200.20 | 23 | 0 | 100.0 | 0.522 |
| 1.10.1740.160 | 24 | 0 | 100.0 | 0.598 |
| 1.10.1840.10 | 40 | 0 | 100.0 | 0.481 |
| 1.10.2000.10 | 24 | 0 | 100.0 | 0.471 |
| 1.10.287.3490 | 13 | 0 | 100.0 | 0.334 |
| 1.10.300.10 | 15 | 0 | 100.0 | 0.543 |
| 1.10.3230.30 | 13 | 0 | 100.0 | 0.296 |
| 1.10.3380.10 | 13 | 0 | 100.0 | 0.374 |
| 1.20.120.720 | 16 | 0 | 100.0 | 0.540 |
| 1.20.120.80 | 11 | 0 | 100.0 | 0.492 |
| 1.20.120.880 | 24 | 0 | 100.0 | 0.468 |
| 1.20.142.10 | 16 | 0 | 100.0 | 0.549 |
| 1.20.5.3310 | 13 | 0 | 100.0 | 0.449 |
| 1.20.5.3790 | 17 | 0 | 100.0 | 0.394 |
| 1.20.930.10 | 14 | 0 | 100.0 | 0.439 |
| 2.140.10.10 | 22 | 0 | 100.0 | 0.473 |
| 2.30.26.10 | 15 | 0 | 100.0 | 0.382 |
| 2.30.31.40 | 12 | 12 | 0.0 | 0.186 |
| 2.40.128.190 | 38 | 0 | 100.0 | 0.321 |
| 2.40.128.260 | 17 | 0 | 100.0 | 0.548 |
| 2.60.30.10 | 146 | 0 | 100.0 | 0.422 |
| 2.60.40.1210 | 12 | 0 | 100.0 | 0.570 |
| 2.60.40.1490 | 15 | 0 | 100.0 | 0.404 |
| 2.60.40.1520 | 15 | 0 | 100.0 | 0.502 |
| 2.60.40.180 | 89 | 0 | 100.0 | 0.494 |
| 2.60.40.1940 | 11 | 0 | 100.0 | 0.275 |
| 2.60.40.2470 | 11 | 0 | 100.0 | 0.496 |
| 2.60.40.690 | 15 | 0 | 100.0 | 0.537 |
| 2.60.40.780 | 124 | 0 | 100.0 | 0.534 |
| 2.70.100.10 | 143 | 0 | 100.0 | 0.471 |
| 3.10.450.360 | 15 | 7 | 53.3 | 0.226 |
| 3.20.16.10 | 22 | 0 | 100.0 | 0.549 |
| 3.30.1280.10 | 14 | 0 | 100.0 | 0.478 |
| 3.30.1350.10 | 15 | 0 | 100.0 | 0.443 |
| 3.30.1360.80 | 24 | 0 | 100.0 | 0.528 |
| 3.30.1740.10 | 15 | 0 | 100.0 | 0.531 |
| 3.30.1740.20 | 12 | 0 | 100.0 | 0.396 |
| 3.30.420.230 | 15 | 0 | 100.0 | 0.424 |

*Continued on next page*

Table S10: **Per-superfamily rejection in the frozen 50-H absent-reference cohort** (continued).

| CATH H | Domains | False accepts | Rejected (%) | Median $d$ |
|---|---|---|---|---|
| 3.30.800.10 | 11 | 0 | 100.0 | 0.350 |
| 3.40.1210.10 | 18 | 0 | 100.0 | 0.536 |
| 3.40.30.120 | 14 | 0 | 100.0 | 0.352 |
| 3.40.50.10310 | 21 | 0 | 100.0 | 0.459 |
| 3.40.50.12240 | 12 | 0 | 100.0 | 0.480 |
| 3.40.50.9100 | 24 | 0 | 100.0 | 0.451 |
| 3.40.532.10 | 16 | 0 | 100.0 | 0.465 |
| 3.40.630.170 | 15 | 0 | 100.0 | 0.411 |
| 3.90.170.10 | 14 | 0 | 100.0 | 0.539 |
| 3.90.330.10 | 119 | 0 | 100.0 | 0.431 |
| 3.90.340.10 | 29 | 0 | 100.0 | 0.481 |

### Supplementary Methods

#### Result provenance and comparability

All headline numbers use protocol `cath_s20_remote_v2`: the sequence-filtered S20 split (116,301 / 514 / 1,028 train/validation/test domains; 5,659 training superfamilies) and ProstT5 AA‖3Di embeddings from the 11,131,930-row concat store. Main-text Table 1, Figures 2–4 and Supplementary Tables S1, S4, S7 and S10 are scored on that roster under CATH-train lookup. Supplementary Table S2 is the three-seed S20 AA‖3Di audit of the production CATH+TED heads. Supplementary Table S3 isolates frozen ProstT5, a CATH-only CCL head and the production CATH+TED head on the same S20 AA‖3Di inputs (seed 40). Development results from dual-filtered or S40 production encoders are not included here.

#### Implemented center-contrastive objective

Let a mini-batch contain $N$ projected embeddings $\mathbf{z}_i \in \mathbb{R}^d$ with H-level labels $y_i \in \{1, \ldots, K\}$. The projection head L2-normalizes every embedding. The loss maintains one raw learnable parameter vector per class and uses its differentiably normalized form $\mathbf{c}_j$ in each forward pass. Define

$$u_{ij} = s\left(\mathbf{z}_i^\top \mathbf{c}_j - m\,\mathbb{1}[j = y_i]\right), \qquad p_{ij} = \frac{\exp(u_{ij})}{\sum_{k=1}^{K} \exp(u_{ik})}. \tag{1}$$

Thus the additive cosine margin is subtracted only from the target logit and is multiplied by the same scale $s$ as the cosine similarities. With label smoothing $\varepsilon$, the target distribution implemented in the code is

$$q_{ij} = \begin{cases} 1 - \varepsilon, & j = y_i, \\ \varepsilon/(K-1), & j \neq y_i. \end{cases} \tag{2}$$

The batch objective is

$$\mathcal{L}_{\text{CCL}} = \frac{1}{N} \sum_{i=1}^{N} \left[ -\sum_{j=1}^{K} q_{ij} \log p_{ij} + \lambda \left\| \mathbf{z}_i - \mathbf{c}_{y_i} \right\|_2^2 \right]. \tag{3}$$

For unit vectors the second term is $\lambda(2 - 2\mathbf{z}_i^\top \mathbf{c}_{y_i})$. When $\varepsilon = 0$, Equation 3 is the large-margin center-contrastive objective of Cai et al. plus the additive constant $2\lambda$ per sample that is omitted in their algebraically simplified Equation 5; the constant changes the reported loss value but not its gradients. Label smoothing is a stated implementation extension and applies only to the normalized-softmax term; the center-attraction term always uses the hard H-level label.

For CATH's long tail, a sampled singleton class still contributes both a target-versus-all-centers softmax signal and a center-attraction signal. The loss forms an $N \times K$ domain–center similarity matrix and therefore does not require a second member of the same class in the mini-batch. The sampler selects H classes uniformly without replacement and draws at most two domains from each selected class, giving 1,024 domains per batch and 112 batches per epoch. In the CATH S20 training roster, 1,679 of 5,659 H classes are singletons, 2,621 (46.3%) contain at most two domains and the largest class contains 10,132. Uniform class sampling prevents the largest superfamilies from contributing in direct proportion to their abundance. By contrast, batch-hard triplet training forms an $N \times N$ domain–domain distance matrix and can construct an H-level anchor–positive pair only when two members of that superfamily occur in the batch. Training centers are optimizer parameters and are not used directly for the reported retrieval metrics. Nearest-centroid inference instead averages the projected CATH training embeddings belonging to each H class and L2-normalizes those empirical means.

The production settings are $s$=32, $m$=0.3, $\lambda$=1.0 and $\varepsilon$=0.1 (Supplementary Table S5). The code promotes embeddings to fp32 before forming the similarity expression within bf16 mixed-precision training; the reported checkpoints therefore do not use a fully fp32 loss matrix.

#### Training-set augmentation versus CATH-only CCL

Main-text Table 1 reports production ContrasTED under CATH+TED training on sequence-filtered TED domains. Supplementary Table S3 places the seed-40 production AA‖3Di head beside raw ProstT5-AA‖3Di and a matched CATH-only CCL encoder, all under CATH-train lookup on the S20 test roster ($n$=1028). Frozen ProstT5-AA‖3Di reaches 81.9% 1-NN / 73.2% centroid. The CATH-only CCL head reaches 87.8% / 86.4% (+5.9 / +13.2 percentage points). The production CATH+TED seed-40 head reaches 92.3% / 90.7%, a further +4.5 / +4.3 points over CATH-only. TED rows are present in the concat store but were never sampled for the CATH-only run, because `train.fasta` is CATH-only.

#### TED augmentation labels and held-out-set filtering

The augmentation roster originates from the H-labelled portion of TED built from AlphaFold DB v4 domains described in the main manuscript. TED transferred these H labels from CATH using Foldseek structural search. The 11,131,930 retained domains were not required to have an independent profile-HMM assignment agreeing with the Foldseek-derived H label; accordingly, the manuscript treats them as automated or pseudo-labelled training examples. The independent TED Gold NR S20 evaluation requires agreement between Foldseek and profile-HMM H labels.

Sequence filtering removes TED domains at $\geq$ 20% sequence identity to any S20 validation or test query. It excludes sequence-near examples but does not remove structurally similar, sequence-remote domains. No TED domain enters the CATH-train lookup set used for Table 1. Split manifests, retained TED identifiers, search outputs and production training configurations are included with the release.

#### Seed stability of CATH+TED S20 encoders

Production ContrasTED heads are trained on the CATH+TED mixture with a CATH-only validation 1-NN reference set (main-text Methods). To assess seed dependence, we retrained the S20 AA‖3Di arm at seeds 40, 41 and 42 under the locked hyperparameter recipe and scored test H-level accuracy ($\mathrm{Acc_H}$) with the CATH-train lookup (Supplementary Table S2). 1-NN accuracy is 92.9% ± 0.5% and centroid accuracy is 91.4% ± 0.7%. Main-text Table 1 reports those three-seed means; main-text Figures 2–4 use seed 40. The encoder was trained on the S20 concat store (11,131,930 embeddings; all 116,301 / 514 / 1,028 S20 train/validation/test domains resolved).

#### HH-suite3 and TEDLH profile subsets

Table 1 reports HH-suite3 against the train-only TEDLH subset: profiles whose CATH seed domain belongs to the S20 training split [5, 1]. That subset prevents a query from matching a profile seeded on itself and removes profiles seeded on domains withheld from training by the S20 identity filter, matching the CATH-train lookup used by the other methods. The train-only TEDLH subset spans 4,664 of the 5,659 S20 training superfamilies, so 959 of 1,028 test queries have a correct answer available (Supplementary Table S8). Lite and full libraries contain test-domain seeds and are leakage diagnostics, not Table 1 controls (Supplementary Table S9).

#### TED T→H threshold selection

For each checkpoint, we selected the maximum observed validation cosine distance retaining at least 99% point precision among accepted assignments, then fixed that threshold before test evaluation. Applied to the production CATH+TED S20 seed-40 encoder with nearest-centroid retrieval, this rule gives $d = 0.24048$, 99.14% validation precision at 68.09% coverage, and 98.48% locked test precision at 64.01% coverage (658 accepted of 1,028; Supplementary Table S7). This is the threshold used for TED T-hit assignment in main-text Figure 4c–g. Under 1-NN retrieval on the same encoder, achieving $\geq$ 99% validation precision required $d = 0.17817$, yielding 99.06% validation precision at 62.26% coverage and 98.66% test precision at 58.27% coverage (599 accepted of 1,028, 591 correct). Centroid retrieval thus yielded 57 additional correct remote-homology assignments under the 99% precision target. At a 98% validation precision target, centroid retrieval gave 78.11% test coverage (98.01% precision; $d = 0.31691$) versus 77.92% for 1-NN (97.88% precision; $d = 0.26948$).

The 50-superfamily holdout uses a separately trained S20 seed-40 encoder and therefore has its own validation-selected threshold, $d = 0.22579$. At this point, 329/332 accepted validation predictions were correct (99.10% precision; 64.59% coverage). Applying the threshold unchanged gave 98.45% precision at 62.84% coverage on the known-superfamily S20 test set and 98.64% rejection of 1,401 domains from the 50 superfamilies absent from training and lookup. The held-out-superfamily and TED Gold cohorts did not contribute to threshold selection.

#### Output-dimensionality control

We varied the projected dimension over $\{16, 32, 64, 128, 256, 512\}$ using CATH-only AA‖3Di training on the S20 split and the locked center-contrastive hyperparameters (Figure S1). Accuracy rises steeply through 128 dimensions. Some seeds gain modestly at 256 or 512 dimensions, but this exploratory sweep was not repeated after adding sequence-filtered TED training. We therefore retain the pre-specified 128-dimensional production representation, which also matches ProtTucker's output width and provides a 16-fold compression from the 2048-dimensional input.

##### Encoder-layer leave-one-out probe

To locate where remote-homology signal emerges in frozen pLM representations, we mean-pooled every encoder hidden state in a single forward pass and scored leave-one-out (LOO) 1-NN homologous superfamily (H) accuracy at each layer. The probe used the official CATH c1–c3 non-redundant S20 and S40 test rosters after dropping H-singleton superfamilies ($n$=11,057 and 30,480 domains). Each query was classified by cosine 1-NN against all other domains in the split, excluding itself. Figure S2 pools the two splits with domain-count weighting and compares ProstT5 with Biohub ESMC models at matched scales [3]. Figure S3 repeats the probe for ProstT5-AA (<AA2fold>), ProstT5-3Di (<fold2AA>) and concatenated AA‖3Di embeddings on each split separately. These probes diagnose frozen encoder geometry; they do not use the ContrasTED projection head or CATH-train lookup set from Table 1.

#### Supplementary Figures

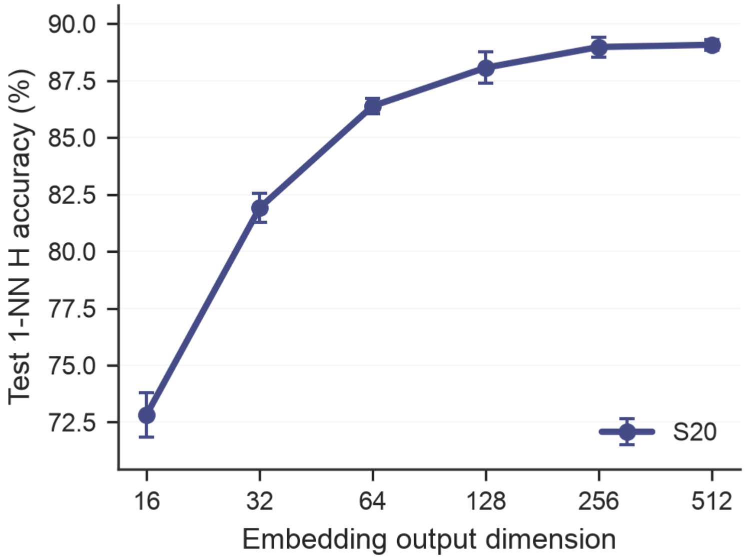


Figure S1: **Output-dimensionality control on the S20 split.** Test 1-NN H accuracy as a function of projected dimension for CATH-only center-contrastive training on ProstT5 AA‖3Di embeddings (lr 0.012, $s$=32, $m$=0.3, $\lambda$=1.0, seeds 40–42, $n$=1028). Points are mean ± s.d. over the three seeds. The production model uses a 128-dimensional representation. The sweep is a dimensionality control rather than an evaluation of the production CATH+TED encoder.

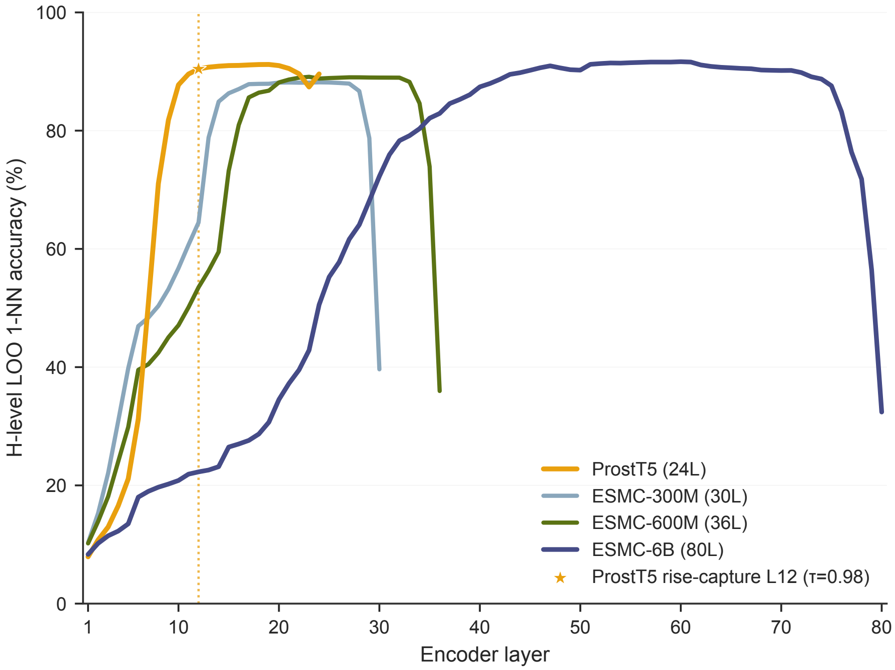


Figure S2: **Encoder-layer remote-homology signal in frozen pLMs.** Domain-weighted mean of S20 and S40 leave-one-out 1-NN homologous-superfamily (H) accuracy versus encoder layer for ProstT5 and Biohub ESMC models (300M / 600M / 6B) [3]. Each series stops at that model's final layer. The vertical marker shows the earliest ProstT5 layer whose accuracy captures at least 98% of the split-wise rise to the peak on both benchmarks (layer 12; $\tau$=0.98). Main-text embeddings use the final ProstT5 layer.

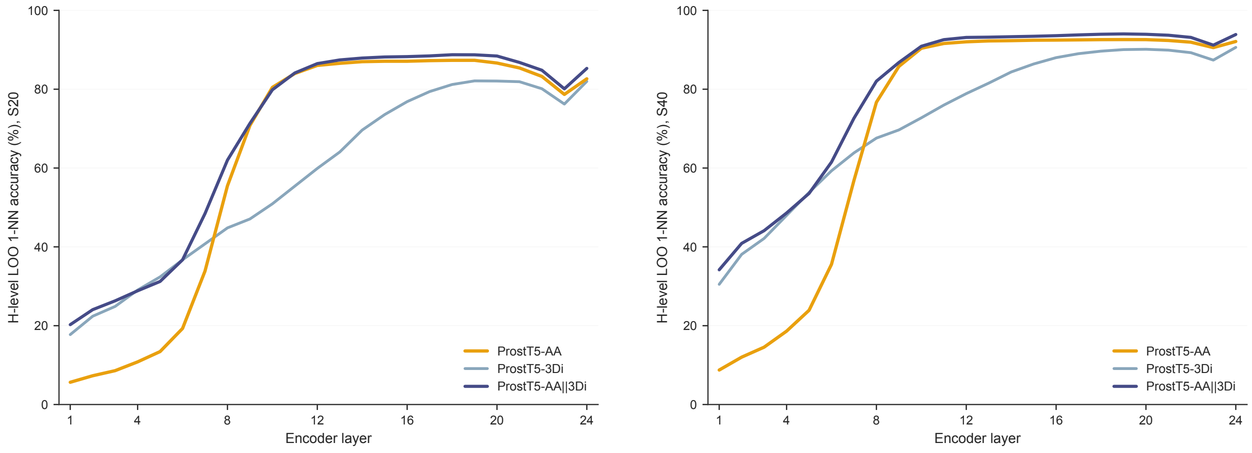


**a** S20 ($n$=11,057) **b** S40 ($n$=30,480)

Figure S3: **ProstT5 modality decomposition across encoder layers.** Leave-one-out 1-NN H accuracy versus encoder layer for mean-pooled ProstT5-AA, ProstT5-3Di and concatenated AA‖3Di embeddings on the (**a**) S20 and (**b**) S40 remote-homology benchmarks. Concatenation improves accuracy throughout the stack; the production pipeline uses final-layer embeddings from both encoder passes.

[2] Matthijs Douze, Alexandr Guzhva, Chengqi Deng, Jeff Johnson, Gergely Szilvasy, Pierre-Emmanuel Mazaré, Maria Lomeli, Lucas Hosseini, and Hervé Jégou. The Faiss library. *arXiv preprint arXiv:2401.08281*, 2024.

[3] EvolutionaryScale. ESM Cambrian: Revealing the mysteries of proteins with unsupervised learning. `https://www.evolutionaryscale.ai/blog/esm-cambrian`, 2024. Biohub ESMC 300M/600M/6B models.

[4] Ilya Loshchilov and Frank Hutter. Decoupled weight decay regularization. In *International Conference on Learning Representations*, 2019.

[5] Martin Steinegger, Markus Meier, Milot Mirdita, Harald Vöhringer, Stephan J. Haunsberger, and Johannes Söding. HH-suite3 for fast remote homology detection and deep protein annotation. *BMC Bioinformatics*, 20(1):473, 2019.
